# An in silico genome-wide screen for circadian clock strength in human samples

**DOI:** 10.1101/2022.05.10.491250

**Authors:** Gang Wu, Marc D. Ruben, Lauren J. Francey, Yin Yeng Lee, Ron C. Anafi, John B. Hogenesch

## Abstract

Years of time-series gene expression studies have built a strong understanding of clock-controlled pathways across species. However, comparatively little is known about how ‘non-clock’ pathways influence clock function. We developed a new computational approach to explore candidate pathways coupled to the clock in human tissues. This method, termed LTM, is an in silico screen to infer genetic influences on circadian clock function. LTM uses natural variation in gene expression in human data and directly links gene expression variation to clock strength independent of longitudinal data. We applied LTM to three human skin and one melanoma datasets and found that the cell cycle is the top candidate clock-coupled pathway in healthy skin. In addition, we applied LTM to thousands of tumor samples from 11 cancer types in the TCGA database and found that extracellular matrix organization-related pathways are tightly associated with the clock strength in humans. Further analysis shows that clock strength in tumor samples are correlated with the proportion of cancer-associated fibroblasts and endothelial cells. Therefore, we show both the power of LTM in predicting clock-coupled pathways and classify factors associated with clock strength in human tissues. LTM is available on GitHub to facilitate its use.

## Introduction

The circadian clock network controls ~24 h rhythms at the cell, tissue, and organismal level. The core network is composed of circadian activators (e.g., BMAL1/CLOCK, RORs and DBP) that bind to Ebox, RORE, and D-box elements to activate transcription of the circadian repressors (e.g., PERs/CRYs, REV-ERBs, NFIL3, DECs and CHRONO), and other clock regulators (e.g., CSNK1D/E and FBXL3/21) (Takahashi, 2017). This feedback mechanism regulates the expression of hundreds to thousands of genes in the mammalian genome (Zhang et al., 2014). Importantly, increasing evidence suggests that the clock network and its time-keeping properties are influenced by other pathways. For example, hepatocyte-specific ablation of *Smo,* a key component of Hedgehog signaling, decreased the amplitude of clock genes (*Bmal1, Clock* and *Nr1d2*) in mouse liver (Marbach-Breitrück et al., 2019). We need a strong understanding of clock-coupled pathways in human tissues to better appreciate the links between disease and clock function.

Previously, efforts were made to explore clock-coupled pathways in human U2OS cells by genome-wide knockdown using small interfering RNAs (siRNAs) (Zhang et al., 2009). U2OS cells are widely used circadian reporter model and show robust oscillations of luciferase expression with a period length of 24 h after synchronization. Hundreds of genes had strong circadian phenotypes, including period length changes and/or increases in amplitude in response to knockdown (Zhang et al., 2009). Pathway analysis revealed that insulin and Hedgehog signaling are coupled to the circadian clock in U2OS cells. However, an important limitation of the U2OS circadian reporter cells is that they do not model mammalian circadian systems at the tissue level.

Computational approaches can be used to explore clock-coupled pathways in human tissues. Indeed, the machine learning-based CYCLOPS was developed to identify cycling genes in a human tissue without the need for collection time of day (Anafi et al., 2017). CYCLOPS was applied to multiple human tissues (Ruben et al., 2018; Wu et al., 2018). However, it has limitations, including optimal seed gene list selection (Ruben et al., 2018; Wu et al., 2020) and challenges in evaluating the quality of ordering without known time-stamped samples (Wu et al., 2018). These issues prevented our ability to order many publicly available datasets. Perhaps most importantly, and a general limitation of computational approaches in this field, is that they focus on identifying cycling genes. However, a gene that is cycling may not directly impact clock function. Conversely, a gene that is coupled to the clock network, but is not cycling may directly impact clock function.

We developed a new computational approach to explore candidate pathways coupled to the clock in human population samples without time information. This method, named LTM, is an in silico screen mimicking genome-wide knockdown experiments done in U2OS cells. As opposed to genetic manipulation, we use the natural variation in expression levels of genes in human population datasets to infer genes’ influence on circadian clock function. The idea is that genes whose expression correlates with circadian clock function are clock-coupled candidates. The power of this approach is that it (i) directly links expression variability of a gene to clock function, (ii) does not depend on the longitudinal order of samples by timestamps or machine learning inference, and (iii) predicts pathways coupled to the clock network and/or the factors correlated with clock strength in a large scale dataset.

We applied LTM to human population datasets to identify clock-coupled pathways in skin. We found that the cell cycle is the top candidate clock-coupled pathway in healthy skin, but not in skin tumors. In addition, we applied LTM to thousands of tumor samples in the TCGA database and found that extracellular matrix organization-related pathways are tightly associated with the clock strength in human tumors. We released the LTM on GitHub to facilitate its use.

## Results

### In silico screen for genes associated with circadian clock strength in population data

Our prior work showed that Mantel’s zstat (Wu et al., 2020, 2018) and nCV (Wu et al., 2021) provide reliable measures of clock strength (Fig. S1 A-C) from population samples. Using these two measures of clock strength, we developed a new method, LTM, to screen for genes associated with clock strength in population data. The principle behind LTM is that genes whose variation in expression correlates with variation in clock strength are potentially coupled to the molecular clock.

We used human epidermis data (18,076 genes and 298 samples from 239 individual human donors) (Wu et al., 2018) to explain our strategy (Fig. 1). The schematic focuses on one example gene, *CCNA2,* to demonstrate the major steps of LTM. *CCNA2* was chosen because it is one of the top gene by LTM score. First, we separated each of the 298 samples into one of four quantile groups (Q1, Q2, Q3 and Q4) based on their expression level of *CCNA2.* Next, we calculated the correlation value between the mean expression of *CCNA2* and clock strength per quantile group, resulting in the terms R(nCV) and R(zstat). The mean value of R(zstat) and R(nCV) is defined by LTMpre. Then we repeated this same process twice more, screening at 7 and 10 quantile groups, resulting in two additional LTMpre values for *CCNA2.* Finally, we computed an LTMori and LTMabs value for *CCNA2.* LTMori is the average of multiple LTMpre values. A positive and negative LTMori value means that samples in a quantile group with higher (eg., *CCNA2)* and lower expression (eg., *ROR1)* of this gene has a stronger clock, respectively. LTMabs is the absolute value of LTMori. The LTMabs value makes it easier to explore clock-coupled pathways given that up- or down-regulated genes in the same pathway may affect the circadian clock similarly. In sum, genes with higher LTMabs values represent a stronger correlation between expression level and clock strength. Next, we applied LTM to human population skin samples collected from healthy subjects in multiple sources.

**Figure 1.**
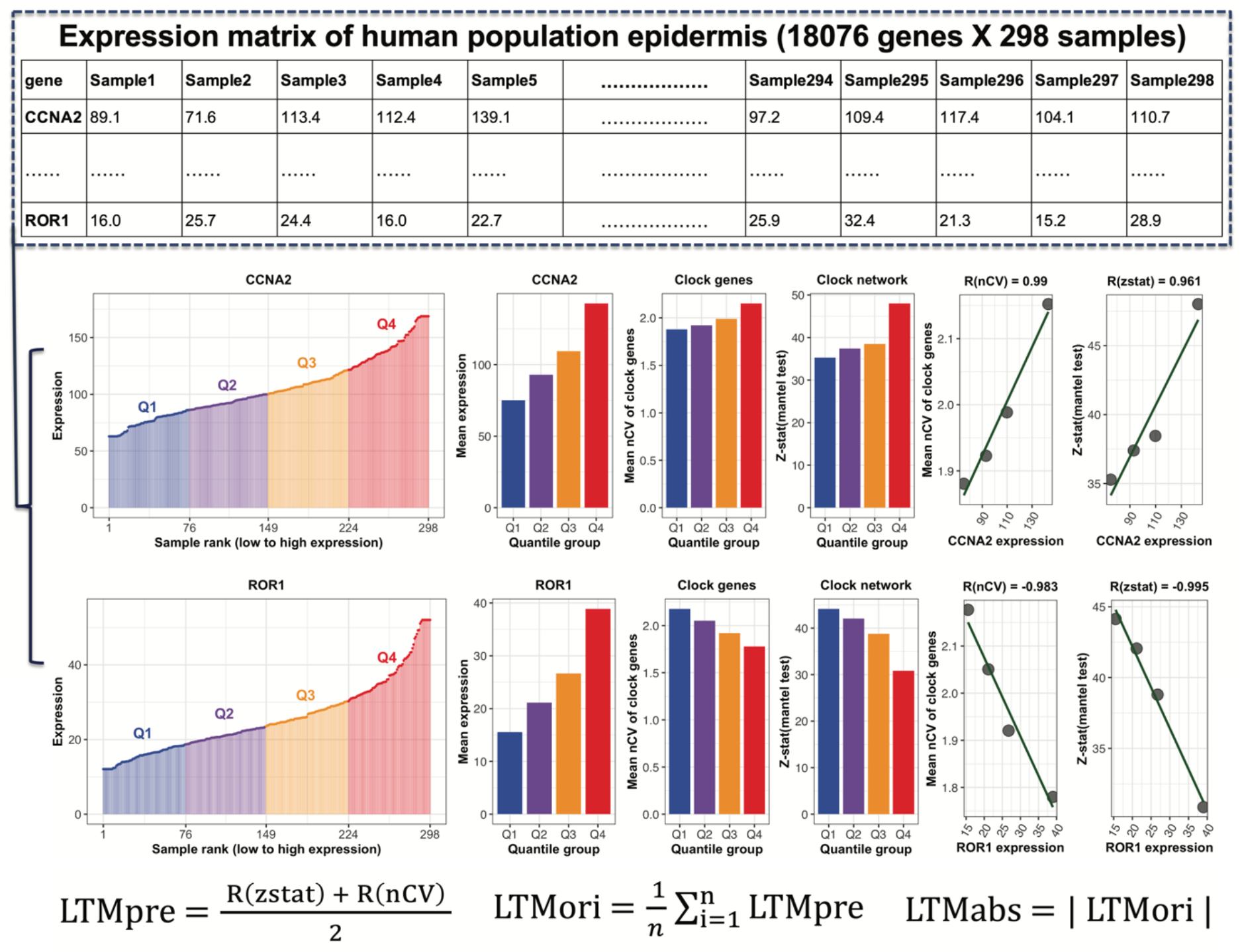
The schematic diagram of LTM strategy. The expression matrix contains 18,076 genes (one gene per row) and 298 samples (one sample per column). All 298 samples are ranked by the expression level of each gene (eg., *CCNA2*) and separated into four quantile groups. There are 75 (Q1 and Q3) or 74 (Q2 and Q4) samples in each quantile group. The mean expression value of all samples in each quartile group is calculated, as indicated by barplot. Two clock network properties were calculated for each quantile group. The first property is clock robustness, that is indicated by the mean nCV of 17 clock and clock-associated genes in Q1 to Q4 separated by *CCNA2’s* expression. The second property is phase conservation at network level, which is indicated by Mantel’s zstat value. The correlation matrix of 17 clock and clock-associated genes in mouse circadian atlas and in each quantile group serves as the reference and query matrix, respectively. The zstat value of the Mantel’s test indicates the similarity between a query matrix and the reference matrix. Higher mean nCV value and Mantel’s zstat value means a stronger clock. The correlation value between mean nCV or Mantel’s zstat value and mean expression of *CCNA2 (ROR1)* from Q1 to Q4 is named as R(nCV) or R(zstat). The LTMpre is defined as the mean value of R(nCV) and R(zstat) of *CCNA2 (ROR1)*. The LTMori is averaged LTMpre values for the same gene. The absolute value of LTMori is defined as the LTMabs.

### Meta-analysis of three human population skin datasets using LTM

We observed inter-individual variation in skin clock strength in our prior study of 20 subjects sampled longitudinally over four timepoints (Wu et al., 2020). Therefore, we expected a range of clock strengths among single skin samples collected from hundreds of human subjects. On the other hand, expression levels of thousands of genes show variation among human skin samples (Kimball et al., 2018). LTM searches for strong correlations between gene expression levels and clock strength, which we applied to three skin datasets: (i) epidermis samples from 239 subjects (PG), (ii) sunexposed whole skin samples from 601 subjects (GTExSE), and (iii) non-sun exposed whole skin samples from 479 subjects (GTExNSE). The distributions of LTMabs values were similar in three skin datasets (Fig. 2A). To control for dataset-specific bias, we computed an integrated-LTMabs value to identify genes most strongly correlated with clock strength across datasets. For example, *IPO13* ranks among the top integrated-LTMabs values indicating a strong linear correlation between its expression and mean nCV (R(nCV) > 0.87 in all three datasets) or Mantel zstat value (R(zstat) > 0.86 in two datasets) (Fig. 2B). *IPO13* encodes a member of the importin-beta family of nuclear transport proteins. Importin-beta genes have been implicated in circadian clock protein transport and function in human cells (Lee et al., 2015; Zhang et al., 2009). LTM identifies *IPO13* as an importin-beta gene with a particularly strong association to circadian clock function in human skin.

**Figure 2.**
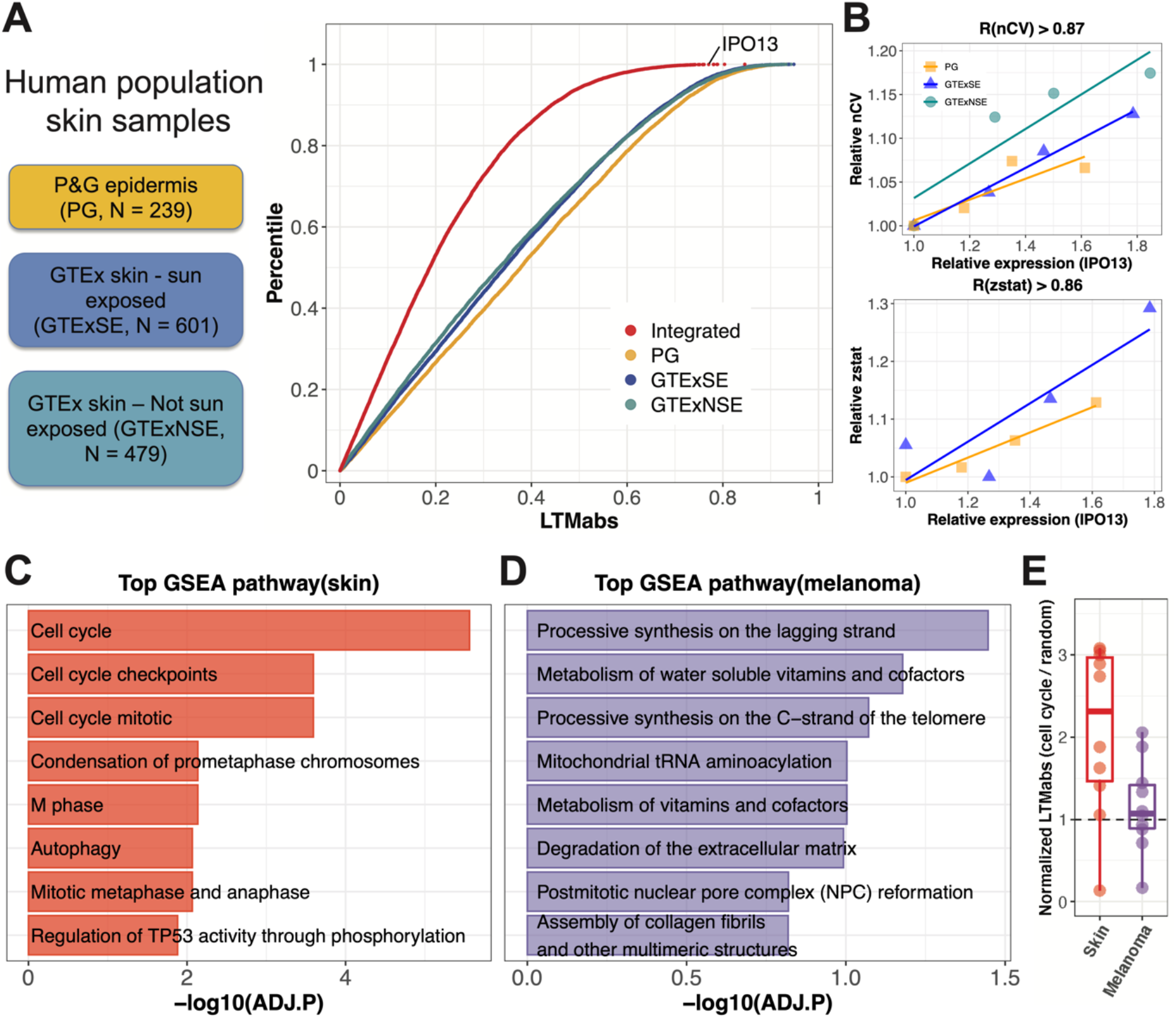
Circadian clock-coupled pathways screened by LTM in human population skin samples. (A) LTM is used to screen three groups of human population skin samples, including human epidermis samples from 239 subjects (PG), sun exposed whole skin samples from 601 subjects (GTExSE), and not sun exposed whole skin samples from 479 subjects (GTExNSE). The distribution of LTMabs for PG, GTExSE and GTExNSE is indicated by orange, blue and cyan-blue points respectively. Each red point represents LTMabs of a gene by integrating three datasets. A gene with a large LTMabs (eg. *IPO13*) means its expression level is strongly correlated with clock strength in skin. (B) The expression level of *IPO13* is linearly correlated with mean nCV of clock genes and Mantel zstat in skin datasets. (C) Top 8 enriched pathways from GSEA analysis on all genes ranked by integrated LTMabs of three healthy skin datasets. (D) Top 8 enriched pathways from GSEA analysis on all genes ranked by LTMabs of skin cutaneous melanoma in TCGA database. (E) Cell cycle regulators show lower LTMabs values in human skin tumors than healthy skin. Each point represents a cell cycle gene listed in Fig. S3A. Only those cell cycle genes detected in both healthy skin and melanoma datasets were used. Y-axis is the LTMabs value of each cell cycle gene normalized by the LTMabs value of randomly selected genes.

### Cell-cycle gene expression correlates with clock strength in healthy human skin

We performed gene set enrichment analysis (GSEA) of genes ranked by integrated-LTMabs. Cell cycle regulation was among the top significantly enriched pathways (Fig. 2C; ADJ.P < 0.01). Specifically, genes regulating G2/M phase (e.g., *CDK1, CCNB1* and *CCNB2)* have higher integrated-LTMabs values (ie., their expression tightly correlates with clock strength) compared to genes regulating G1/S phase (eg., *CDK4/6, CCND1,* and *CCND2)* (Fig. S2A-B). *CDK1* and cyclin B genes *(CCNB1* and *CCNB2)* are critical components of M-phase-promoting factor (Hara et al., 2012; Nurse, 1990), which is the universal inducer of M-phase in eukaryotic cells. In sum, LTM found a strong correlation between mitotic entry and circadian clock strength in healthy skin samples.

Highly active cell proliferation is common in human tumors. Is mitotic entry coupled to the circadian clock in skin tumors? Using LTM, we screened 472 tumor samples of skin cutaneous melanoma from the TCGA database. Top ranked pathways include processive synthesis on the lagging strand, metabolism of water-soluble vitamins and cofactors, and processive synthesis on the C-strand of the telomere, but none of them were significantly enriched (Fig. 2D; ADJ.P > 0.01). We further compared the LTMabs value of cell cycle regulators between healthy skin and skin tumors. Unlike what we observed in healthy skin samples, the LTMabs values of cell cycle regulators were much lower in tumors (Fig. 2E and Fig. S2C), indicating a weak correlation between their expression levels and clock strength. Together, these results suggest that the coupling between cell cycle and circadian clock observed in healthy skin is disrupted in tumors.

### Pan-cancer analysis using LTM

Next, we asked which pathway is related to the clock strength in tumor samples. We extended LTM screening on tumor samples from other 10 cancer types. Cancer types included urothelial bladder carcinoma (BLCA), breast invasive carcinoma (BRCA), head-neck squamous cell carcinoma (HNSC), clear cell renal cell carcinoma (KIRC), lung adenocarcinoma (LUAD), low grade glioma (LGG), lung squamous cell carcinoma (LUSC), prostate adenocarcinoma (PRAD), stomach adenocarcinoma (STAD), and thyroid carcinoma (THCA). We computed integrated-LTMabs values from total 11 cancer types (Fig. 3A; red point line). *PER1* was among the top ranked genes, with lower expression correlating with weaker clock function in tumors (Fig. 3B; R(nCV) and R(zstat) above 0.5 in 10 of 11 cancer types). *PER1* downregulation has been observed in many tumors, including breast, prostate, glioma and colorectal cancers (Savvidis and Koutsilieris, 2012), giving us confidence in this pancancer analysis. We performed GSEA on all genes ranked by their integrated-LTMabs values. Top significantly enriched pathways include extracellular matrix organization (ECM), degradation of ECM, and ECM proteoglycans (Fig. 3C; ADJ.P < 0.01). These results suggest that ECM related pathways are associated with the clock strength in tumor samples across cancer types.

**Figure 3.**
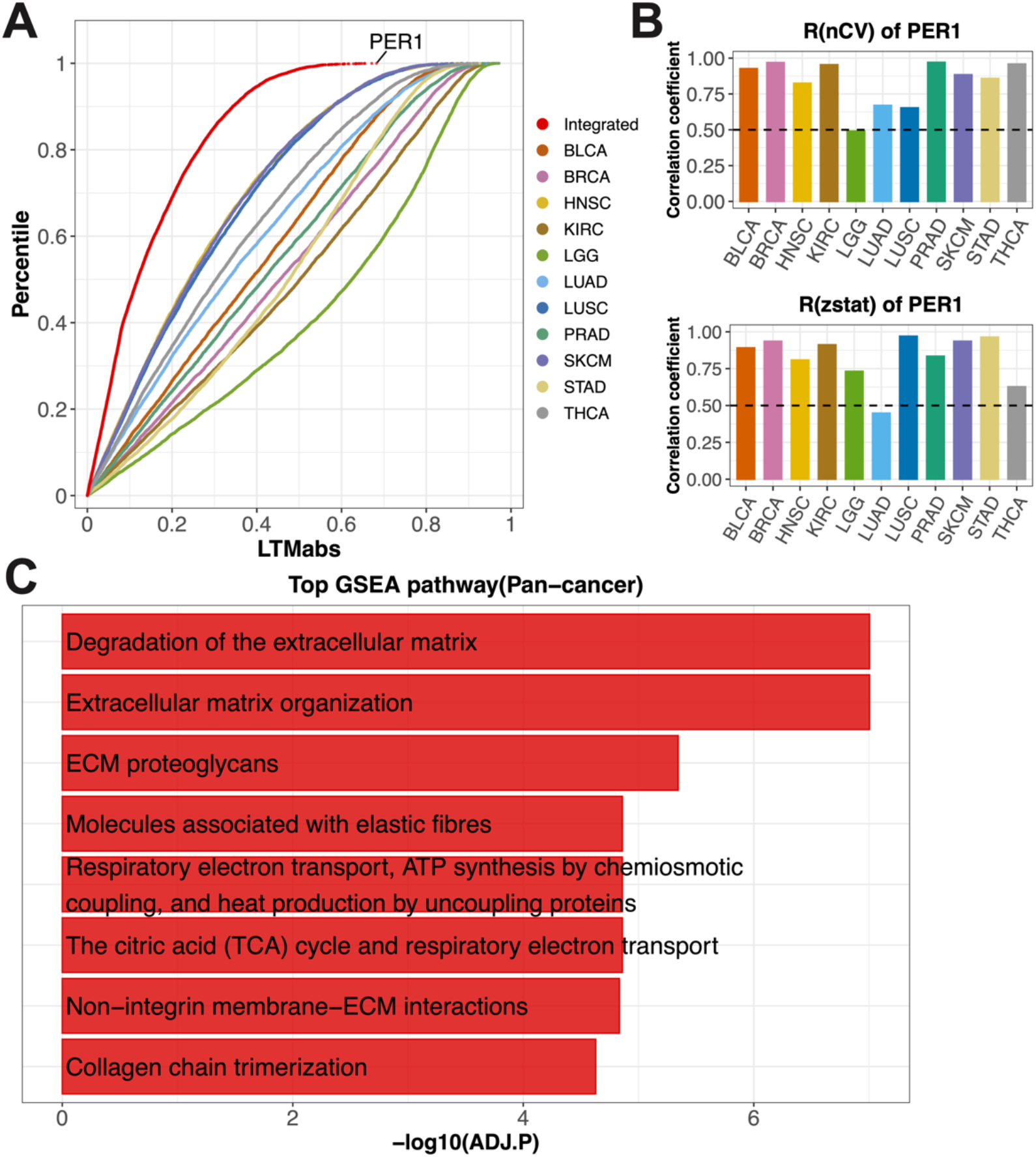
LTM screened genes tightly correlated with clock strength in tumors from 11 cancer types. (A) LTM is used to screen tumor samples of 11 cancer types from the TCGA database, including urothelial bladder carcinoma (BLCA), breast invasive carcinoma (BRCA), head-neck squamous cell carcinoma (HNSC), clear cell renal cell carcinoma (KIRC), lung adenocarcinoma (LUAD), low grade glioma (LGG), lung squamous cell carcinoma (LUSC), prostate adenocarcinoma (PRAD), skin cutaneous melanoma (SKCM), stomach adenocarcinoma (STAD) and thyroid carcinoma (THCA). Different color points indicate the distributions of LTMabs values for different cancer types. Each red point represents the LTMabs value of a gene by integrating 11 datasets. (B) The expression level of *PER1* is positively correlated with mean nCV of clock genes (R(nCV)) and Mantel zstat (R(zstat)) in tumor samples across cancer types. (C) Top 8 enriched pathways from GSEA analysis on all genes ranked by integrated LTMabs values of 11 cancer types.

### Clock strength in tumor samples are correlated with fractions of fibroblast and endothelial cells

Those LTM selected genes in extracellular matrix organization related pathways are primarily collagen genes. We mapped integrated-LTMabs values to the human collagen gene family, which includes 33 genes in 8 subfamilies (Gelse et al., 2003). The collagen genes with high integrated-LTMabs values (LTMabs >= 0.5) are from the fibril-forming, basement membrane, microfibrillar, and multiplexin subfamilies (Fig. 4A). In sum, LTM identified a strong correlation between collagen gene expression and circadian clock strength in tumors.

**Figure 4.**
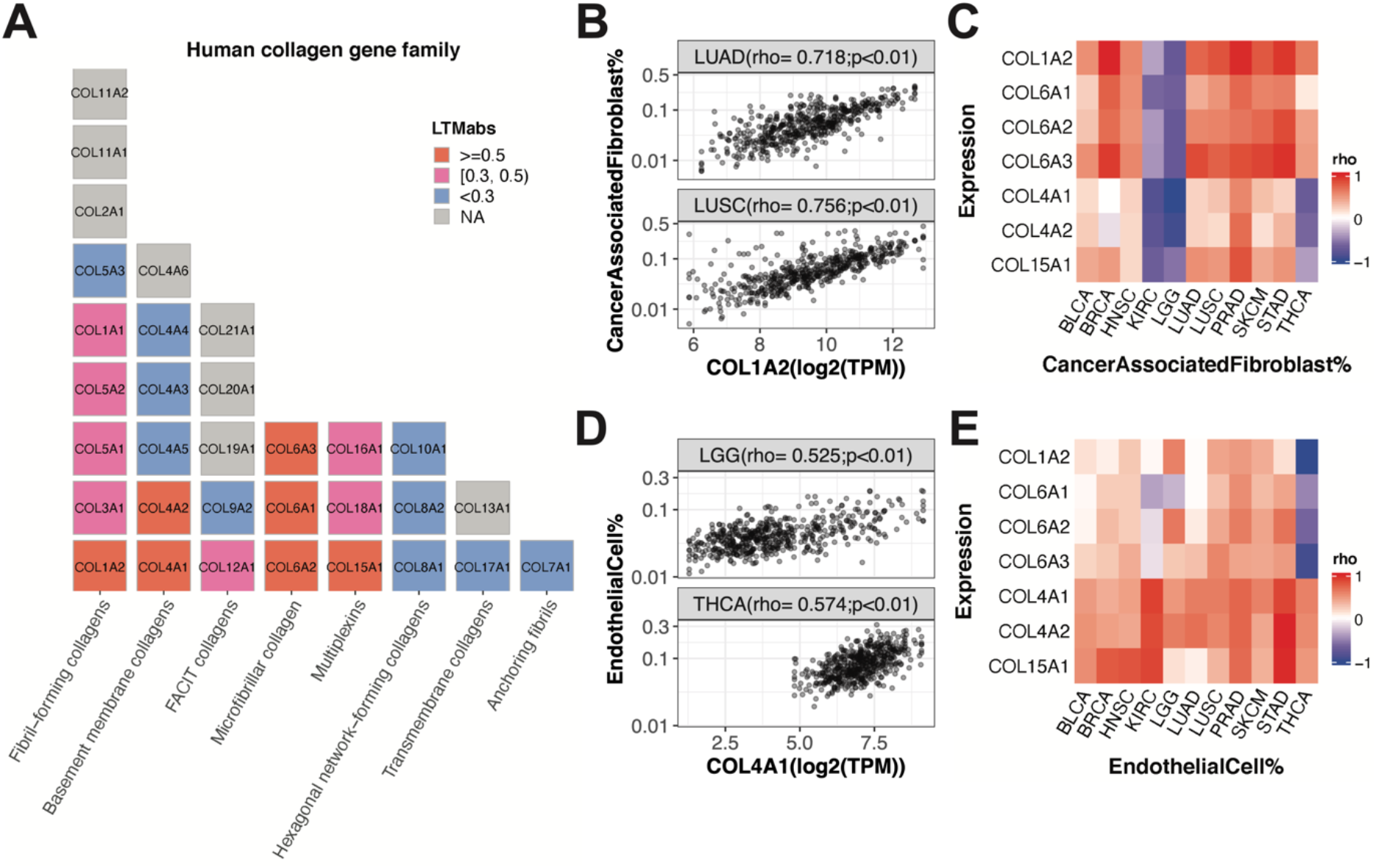
Stromal cell fractions are correlated with expression levels of LTM selected collagen genes. (A) Human collagen genes are listed with a block plot, with one block indicating one gene. Each column represents a collagen category. The LTMabs values are indicated by the block color. (B) The fraction of cancer associated fibroblasts is positively correlated with expression of *COL1A2* in tumor samples from LUAD and LUSC. (C) The fractions of cancer related fibroblast is positively correlated with expression of 7 collagen genes selected by LTM (LTMori >= 0.5) in multiple cancer types. Red and blue colors in the heatmap indicate positive and negative correlation, respectively. (D) The fraction of endothelial cells is positively correlated with expression of *COL4A1* in tumor samples from LGG and THCA. (E) The fractions of the endothelial cell is positively correlated with expression of 7 collagen genes selected by LTM (LTMori >= 0.5) in multiple cancer types. Red and blue colors in the heatmap indicate positive and negative correlation, respectively. The 11 cancer types include urothelial bladder carcinoma (BLCA), breast invasive carcinoma (BRCA), head-neck squamous cell carcinoma (HNSC), clear cell renal cell carcinoma (KIRC), lung adenocarcinoma (LUAD), low grade glioma (LGG), lung squamous cell carcinoma (LUSC), prostate adenocarcinoma (PRAD), skin cutaneous melanoma (SKCM), stomach adenocarcinoma (STAD) and thyroid carcinoma (THCA) from TCGA database.

To understand the basis for this correlation, we asked how tumor samples with low collagen gene expression differ from tumor samples with high collagen gene expression. It is known that Type I collagen genes (e.g., *COL1A1* and *COL1A2)* are highly expressed in human lung fibroblasts (Habermann et al., 2020). This encouraged us to test the correlation between *COL1A2* expression and the proportion of fibroblasts across lung tumor samples. Using EPIC (Racle et al., 2017), we computed the fraction of cancer associated fibroblasts in two lung cancer types, LUAD and LUSC. Indeed, we found a strong positive correlation between *COL1A2* expression and fibroblast proportion in both LUAD and LUSC (Fig. 4B). We extended this analysis to test all the 7 top collagen genes (LTMabs >= 0.5) among 11 cancer types. We found a consistent positive correlation between fibroblast proportions and expression of collagen genes belonging to fibril-forming *(COL1A2)* and microfibrillar *(COL6A1, COL6A1,* and *COL6A3)* subfamilies across 9 cancer types, the exception being clear cell renal cell carcinoma and low grade glioma (Fig. 4C). Overall, this suggests that variation in clock strength between tumors is related to the proportion of fibroblasts in those samples.

On further inspection, we identified two Type-IV collagen genes *(COL4A1* and *COL4A2)* whose expression levels are negatively correlated with fibroblast proportion in thyroid cancer and low grade glioma. It is known that Type-IV collagen plays an important role in endothelial cell adhesion and migration (Herbst et al., 1988). Thus, we measured the correlation between *COL4A1* expression and endothelial cell proportions in thyroid cancer and low grade glioma samples, and found a strong positive correlation (Fig. 4D). The extended analysis of top 7 collagen genes among 11 cancer types show a consistent positive correlation between the fractions of endothelial cells and expression of basement membrane *(COL4A1* and *COL4A2)* and multiplexin *(COL15A1)* collagen subfamily (Fig. 4E). In sum, variation in clock strength among tumor samples is related to the proportion of fibroblasts and endothelial cells. This makes sense considering non-cancer cells including fibroblast and endothelial cells have stronger clocks than cancer cells in the tumor microenvironment.

What causes variation in the relative abundance of fibroblasts and endothelial cells between tumor samples? One possibility is technical bias whereby surgical resection captures more or less surrounding, non-tumor tissue. A second possibility is that tumors vary in cell composition. It is known that the immune cell composition of the tumor microenvironment varies among cancer types and between cancer patients (Ansell and Vonderheide, 2013). If true, perhaps an important question to ask is, do cancer cells surrounded by more fibroblast and endothelial cells have stronger clocks than other cancer cells? More importantly, do clock strength differences in cancer cells impact prognosis? We will leave these questions for later study.

## Discussion

The circadian field has built a strong understanding of clock-controlled pathways from years of time-series gene expression studies across species. However, comparatively little is known about how ‘non-clock’ pathways influence circadian clock function, especially in humans. We developed LTM as a window of insight into clock-coupled pathways in human population scale data.

LTM applied to thousands of healthy human skin samples revealed specific cell cycle genes that are strongly correlated with clock strength. While it is known that the circadian clock regulates progression of the cell cycle (Kowalska et al., 2013; Matsuo et al., 2003), it is far less clear if and how the cell cycle regulates progression of the circadian clock. We found that expression levels of cell cycle genes (e.g.,*CDK1* and *CCNB1)* controlling the G2/M phase are tightly correlated with clock strength in skin samples. What is the benefit of having a strong coupling between circadian clock and the cell cycle? It is known that the circadian clock regulates DNA repair (Gaddameedhi et al., 2011) and the cell cycle (Geyfman et al., 2012) in mouse skin. To minimize DNA damage induced by light exposure, a stronger circadian clock may direct more cells to divide at the right time. More cells dividing at the right time may add additional synchronization signals and further strengthen the skin clock. Therefore, the mutual regulation between the circadian and cell cycles may have evolutionary advantages in mammalian skin.

There is increasing evidence that circadian dysregulation contributes to cancer initiation and progression (Hadadi et al., 2020; Kettner et al., 2016; Papagiannakopoulos et al., 2016). This may be caused by less robust clock gene oscillations in tumor samples (Wu et al., 2021). A weak clock may release its temporal gating of the cell cycle (Kowalska et al., 2013; Matsuo et al., 2003), which may contribute to tumor initiation. The release of circadian gating on cell proliferation may in turn damage the coupling between the circadian clock and the cell cycle. Consistent with this idea, LTM on tumor samples found that expression levels of cell cycle regulators are less correlated with clock strength in skin cancer than in healthy skin.

LTM revealed that the expression levels of collagen genes are tightly correlated with clock strength in tumors from 11 cancer types. Our subsequent analysis identified a strong positive correlation between expression levels of these collagen genes and the proportion of fibroblasts and endothelial cells in tumor samples. However, LTM can not tell us anything about the reason why cell proportions vary among tumor samples. One possibility is that tumor microenvironment differs between cancer patients. For example, cancer cell composition may differ in patients. This may relate to the diversity of molecular clock strength between patients. Patients with a stronger molecular circadian clock may have tumors with higher fractions of fibroblasts and endothelial cells. An alternative explanation is that surgical resection obtained more or less tumor, unintentionally capturing margins. This provides an example that the LTM screened genes in the complex tissues (eg., tumors) may pick other stronger factors (eg., fractions of non-cancer cells) than clock-coupled pathways. Considering multiple factors may influence the clock strength in human population datasets, non-biological factors (eg., batch effects) may also affect the interpretation of LTM screening results. It is important to know the dataset before applying LTM.

LTM has other limitations. First, the genes identified by LTM are based on correlations. The strong correlation is not the direct evidence of causal effect. Therefore, the LTM predicted clock-coupled pathways need experimental validation and/or literature evidence. Second, LTM can not screen genes correlated with the period and phase variation of the circadian clock. Third, ranking samples by the gene expression are strongly influenced by the circadian phase for strong cyclers, which makes it difficult to accurately quantify the clock strength when separating samples by circadian phase. So LTM may be biased to non-cycling genes. To fully explore the power of LTM by the research community and fixing its limitations in the future, we release the LTM method into GitHub.

LTM is an in silico genome-wide screen of genes whose expression correlates with clock strength. This method takes advantage of the natural expression variation of genes in human population datasets. Compared to current rhythmic detection methods, the power of LTM is its ability to directly link expression variability of a gene to clock network properties, allowing for the detection of clock-coupled pathways without requiring known or predicted sampling times. An improved knowledge of clock-coupled pathways is important for the clinical application of circadian medicine. For example, a recent study shows that disruption of the clock gene *BMAL1* impacts insulin sensitivity and liver disease (Jouffe et al., 2022). Another showed that targeting the Hedgehog signaling pathway affects the liver clock (Marbach-Breitrück et al., 2019). Therefore, when applying drugs targeting core clock genes or Hedgehog signaling pathway, the mutual influence needs to be considered. Outside of circadian medicine, there are well-known clinical risks of intervention when mutual regulation of pathways is poorly understood. For example, Vioxx (a COX2 inhibitor) is designed to inhibit pain and inflammation. However, Vioxx also inhibits the cardioprotection intermediates produced by COX2 and increased cardiovascular risk in patients (FitzGerald, 2004). This lead to its withdrawal from the market, just 5 years after FDA approval and after many patients deaths. LTM provides a chance of understanding clock coupling mechanisms at the tissue level, and the translation of circadian medicine in clinical application.

## Materials and Methods

### Datasets

All datasets used in this study are listed in Table S1, including experimental design information, the accession number or download link, and number of total samples for each dataset. The raw RSEM counts downloaded from FireBrowse (http://firebrowse.org/) were transformed to TPM values. The refGenome package (version 1.7.7) was used to parse the human gtf file (human genome version: GRCh38). For a gene with multiple transcript splice isoforms, we selected the longest transcript as the representative gene length.

### Main steps of the LTM strategy

We implemented the LTM strategy into the LTMR package. This R package is released on GitHub (https://github.com/gangwug/LTMR). Major steps of performing LTM analysis include:

1. **Normalize the input expression matrix**. This step can perform quantile normalization across samples, compute the expression profile for each gene if multiple splice-isoforms exist, remove genes with low expression value across samples, and blunt extreme low or high outlier expression values. This step is not essential, but is suggested. LTMR::LTMprep function is designed for this step.
2. **Select genes for LTM analysis**. This step separates samples into four quantile groups (Q1, Q2, Q3, Q4) based on the expression level of each gene. It helps further filter out genes with low expression value in the Q1 group, and/or low fold change between Q4 and Q1. The mean expression value in each quantile group progressively increases from Q1 to Q4. LTMR::LTMcut function is designed for this optional step.
3. **Compute the quantitative measures of circadian clock strength for each quantile group**. LTMR::LTMheat function is designed for this essential step. Given a targeted number (eg., 4) of quantile groups, LTMheat ranks samples by the expression level of each gene and separates them into 4 quantile groups (Q1 to Q4) based on their expression level. LTMheat computes the expression correlation matrix of 17 clock and clock-associated genes (Wu et al., 2018) in each quantile group that serves as the query correlation matrix. The correlation matrix of 17 clock and clock-associated genes of mouse circadian atlas data (Zhang et al., 2014) serves as the reference correlation matrix (Wu et al., 2020). LTMheat applies Mantel’s test to compute the similarity statistical value (Mantel’s zstat value) between the query and reference correlation matrix. LTMheat also computes the nCV values (Wu et al., 2021) of 17 clock and clock-associated genes in each quantile group, and further calculates the mean nCV value of 17 clock and clock-associated genes.
4. **Generate the R(nCV) and R(zstat) for each gene**. LTMR::LTMcook function is designed for this essential step. Given a targeted number (eg., 4) of quantile groups, LTMcook generates a correlation value, R(nCV), between the mean expression value and the mean nCV value of 17 clock and clock-associated genes of 4 quantile groups. LTMcook also generates the correlation value, R(zstat), between the mean expression value and the Mantel’s zstat value of four quantile groups. Both R(nCV) and R(zstat) vary between −1 and 1.
5. **Calculate the LTMabs value for each gene.** LTMR::LTMdish function is designed for this essential step. LTMdish averages the mean value of R(zstat) and R(nCV) as LTMpre, which varies between −1 to 1 for all genes. For the same dataset, multiple LTMpre values will be generated when several targeted numbers of quantile groups are tested. The average of multiple LTMpre values for the same gene is calculated as LTMori, and its absolute value is defined as LTMabs for this gene. The LTMabs of all genes vary between 0 and 1.
6. **Integrate the LTMabs value of multiple datasets.** When multiple datasets are available for the same tissue, this step can integrate multiple LTMori values for each gene and output the integrated value. LTMR::LTMmeta function is designed for this optional step.

### LTM analysis of skin and cancer datasets

The key parameters of LTMR functions used to analyze three skin datasets and 11 cancer datasets are listed in Table S2. Those cell cycle regulation genes that encode cyclin and cyclin-dependent kinase were selected to map the LTMabs value. To compare these selected genes between healthy skin and skin tumors, the LTMabs value of each gene was normalized by the mean LTMabs value of 1000 rounds of randomly sampled genes in each dataset.

### Enrichment analysis of LTM ranked genes in skin and cancer datasets

The integrated LTMabs value of three skin datasets was used to rank the genes. Gene set enrichment analysis (GSEA) of all genes ranked by integrated LTMabs value was performed using the fgsea package (Korotkevich et al., 2021). The ‘c2.cp.v7.1.symbols.gmt’ was downloaded from MsigDB (https://www.gsea-msigdb.org/gsea/msigdb). From this file, we only used those REACTOME pathways with gene numbers between 10 and 1000 as the reference gene set of GSEA. Similar procedures were performed on the GSEA of all genes ranked by LTMabs of melanoma or integrated LTMabs value of 11 cancer types.

### Predicted immune cell fractions of tumor samples from 11 cancer types

The immune cell fractions of tumor samples were predicted with the immunedeconv package (Sturm et al., 2019), using EPIC method (Racle et al., 2017). The predicted fractions of CD4^+^, CD8^+^ T cell, macrophage, natural killer cell and B cell in each tumor sample were used in this study.

## Acknowledgements

We thank members of the Hogenesch lab and Smith lab for valuable discussion and advice on this project. This work was supported by the National Cancer Institute [1R01CA227485-01A1 to Ron Anafi and J.B.H.]; National Institute of Neurological Disorders and Stroke [5R01NS054794-13 to Andrew Liu and J.B.H.]; and the National Heart, Lung and Blood Institute [5R01HL138551-02 to Eric Bittman and J.B.H.].

---

**Figure. S1.**
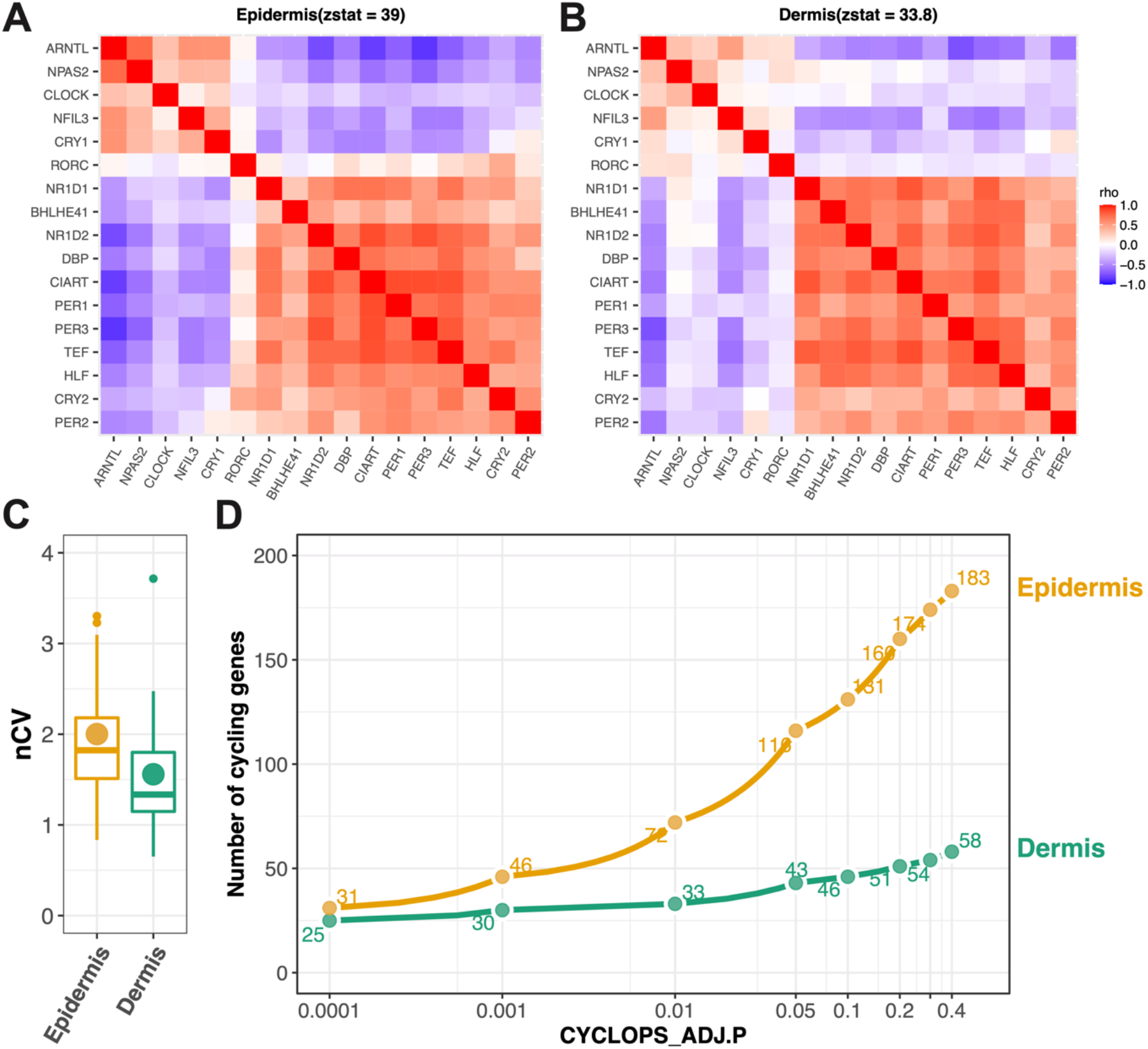
Stronger clock network properties in human epidermis than dermis. Heatmaps of Spearman’s rho for 17 clock and clock-associated genes in human population epidermis (A) and dermis (B) samples. Red and blue indicate positive and negative Spearman’s ρ, respectively. The zstat value is from a Mantel test, using the correlation matrix of clock and clock-associated genes in mouse circadian atlas as the reference. (C) The nCV of 17 clock and clock-associated genes were shown for population epidermis and dermis samples. The point indicates the mean nCV value of 17 genes in each group. The boxes indicate data between the 25th and 75th percentiles with central horizontal lines representing the median values, respectively. The whiskers of boxes show the 5th and 95th percentiles. (D) Number of circadian genes detected in epidermis and dermis samples at a series of ADJ.P values given by CYCLOPS. The data are from Wu G et al., Genome Med, 2020.

**Figure. S2.**
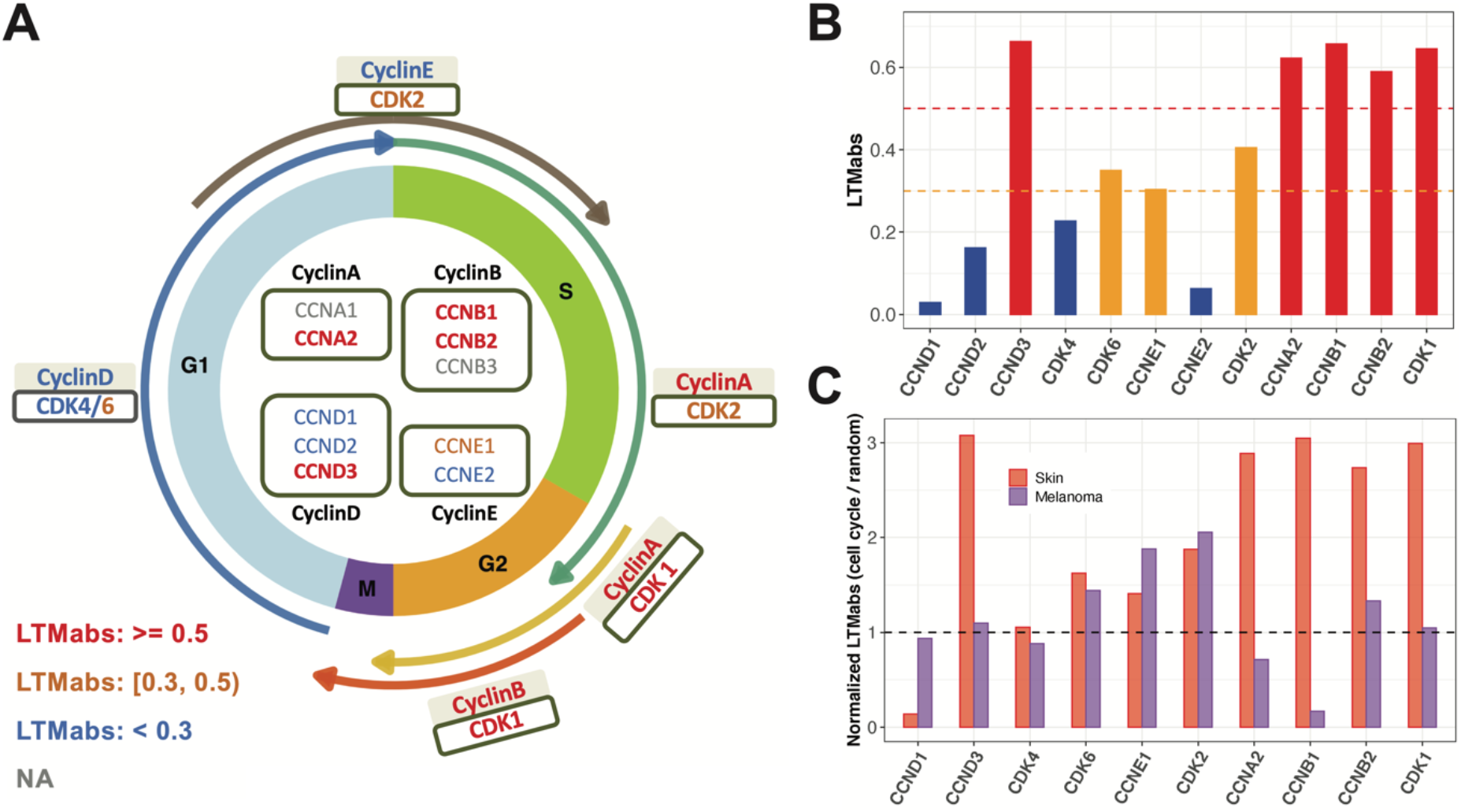
Cell cycle genes with higher LTMabs values screened from human skin datasets regulate G2 and M phase. (A) Genes regulating cell cycle phases are drawn along the circle. The LTMabs value is indicated by text color. (B) The LTMabs value of cell cycle regulators. (C) The normalized LTMabs value of cell cycle genes screened from human skin and melanoma datasets. Only those cell cycle genes detected in both healthy skin and melanoma datasets were used. Y-axis is the LTMabs value of each cell cycle gene normalized by the LTMabs value of randomly selected genes.

**Table S1.** List of datasets used in this study.

| Datasets | #Samples | Experimental design | Reference |
| --- | --- | --- | --- |
| Epidermal skin (PG) | 298 | This dataset includes 79 and 219 samples collected from the longitudinal and population group, respectively. The longitudinal group includes 20 Caucasian male subjects. The population group includes 152 Caucasian female subjects and 67 African-Americans female subjects. | Wu G., et al., 2018, PNAS. GSE112660 |
| Sun exposed whole skin samples (GTExSE) | 601 | The Genotype-Tissue Expression (GTEx) project is to study tissue-specific gene expression and regulation. Samples were collected from 54 non-diseased tissue sites across nearly 1000 individuals. This study uses the version 8 of RNAseq expression profiles in two skin sites. Those skin samples with RIN value less than 6.5 were filtered out in this study. | https://gtexportal.org/home/datasets |
| Not sun exposed whole skin (GTExNSE) | 479 |  |  |
| CYCLOPS ordered epidermal and dermal skin | 489 | 489 matched epidermal and dermal skin samples were selected from 519 CYCLOPS ordered epidermal skin samples and 506 CYCLOPS ordered dermal skin samples. | Wu G., et al., 2020, Genome Medicine. |
| Urothelial bladder carcinoma (BLCA) | 408 |  | http://firebrowse.org/ |
| Breast invasive carcinoma (BRCA) | 1100 | The Cancer Genome Atlas (TCGA) characterized over 20,000 primary cancers and matched normal samples spanning 33 cancer types. This study only used tumor samples from 11 cancer types. Each cancer type contains over 400 tumor samples. |  |
| Head-neck squamous cell carcinoma (HNSC) | 522 |  |  |
| Clear cell renal cell carcinoma (KIRC) | 534 |  |  |
| Low grade glioma (LGG) | 530 |  |  |
| Lung adenocarcinoma (LUAD) | 517 |  |  |
| Lung squamous cell carcinoma (LUSC) | 501 |  |  |
| Prostate adenocarcinoma (PRAD) | 498 |  |  |
| Skin cutaneous melanoma (SKCM) | 472 |  |  |
| Stomach adenocarcinoma (STAD) | 415 |  |  |
| Thyroid carcinoma (THCA) | 509 |  |  |

**Table S2.** The setting of key parameters of LTMR functions in this study.

| Function name | PG | GTE<sub>x</sub>SE | GTE<sub>x</sub>NSE | 11 TCGA cancer types |
| --- | --- | --- | --- | --- |
| LTMprep | quantNorm = TRUE,<br>uniStyle = "mad",<br>removeLowQuant = 0.1,<br>bluntLowQuant = 0.025,<br>bluntHighQuant = 0.975 |  |  | quantNorm = TRUE,<br>uniStyle = "mad",<br>removeLowQuant = 0.25,<br>bluntLowQuant = 0.01,<br>bluntHighQuant = 0.99 |
| LTMcut | minExp = 8 | minExp = 1 | minExp = 1 | minExp = 5 |
| LTMheat | qnum = 4; qnum = 7; qnum = 10 |  |  |  |
| LTMcook | corMethod = "pearson" |  |  |  |
| LTMdish | targetMeasures = c("zmantel", "zncv") |  |  |  |

